# Proteomic insight into human directed evolution of the domesticated chicken *Gallus gallus*

**DOI:** 10.1101/2023.02.07.525900

**Authors:** Carl J Schmidt, Dong Kyun Kim, G Ken Pendarvis, Behnam Abasht, Fiona M McCarthy

## Abstract

Chicken domestication began at least 3,500 years ago for purposes of divination, cockfighting, and food. Prior to industrial scale chicken production, domestication selected larger birds with increased egg production. In the mid-20^th^ century companies began intensive selection with the broiler (meat) industry focusing on improved feed conversion, rapid growth, and breast muscle yield. Here we present proteomic analysis comparing the Ross 708 modern broiler line with the UIUC legacy line. Comparing the breast muscle proteome between modern broilers and legacy lines not selected for these growth traits identifies cellular processes that have responded to human directed evolution. Mass spectrometry was used to identify differences in protein levels in the breast muscle of 6-day old chicks from Modern and Legacy lines. The results highlighted elevated levels of stress proteins, ribosomal proteins, and proteins that participate in the innate immune pathway in the Modern chickens. Furthermore, the comparative analyses indicated differences in the levels of proteins involved in multiple biochemical pathways. In particular, the Modern line had elevated levels of proteins affecting the pentose phosphate pathway, TCA cycle, and fatty acid oxidation and reduced protein levels of the first phase of glycolysis. These analyses provide hypotheses linking the morphometric changes driven by human directed selection to biochemical pathways. The results also have implications for the onset of Wooden Breast disease that arose due to selection for rapid breast muscle growth and is a major problem in the poultry industry.

## Introduction

The chicken has played an important role in human culture and nutrition since its domestication (1, 2). The majority of the modern chicken’s genome was derived from the Red Junglefowl with documented introgression of alleles from the Grey, Ceylon, and Green Junglefowl (3). Early domestication likely led to larger birds that matured quicker and increased egg production in comparison with their wild progenitors (4, 5). The advent of modern agriculture in the early 20th century led to intensive genetic selection for either meat (broiler) or egg (layer) production traits in the chicken. Initial efforts selecting for increased broiler mass led to accumulation of adipose tissue, probably because the selection did not channel the increased level of nutrients to a particular tissue (6). Ultimately, improving broiler production traits led to selection for a combination of larger breast muscle mass, improved feed efficiency and rapid growth.

Comparative studies identifying differences between selected and unselected (legacy) lines provide insight into the impact of evolution of species. Several studies have characterized the differences between modern broilers and legacy lines that have not been subjected to production level human directed selection (7-13). In our work (14-16) we compared the modern Ross 708 broiler line and the legacy University of Illinois, Urbana Campus (UIUC) lines (17, 18). In the legacy line the breast muscle comprises approximately 9% of the body mass, while in the modern broiler this tissue constitutes up to 22% of the body mass (14). Evolving in the tropics, the jungle fowl had little need for long distance flying. The wild chicken is an episodic flier, only needing the ability to fly up into a tree to escape predators and to roost. Like the legacy line, the breast muscle of the Red Jungle Fowl constitutes approximately 9% of its body mass (19). For comparison, the breast muscle of birds capable of more sustained flight averages 17% while that of hummingbirds varies between 25%-30% (20).

The increase in the modern broiler’s breast muscle mass, which almost reaches the hummingbird level, came at the expense of other tissues. For example, the normalized masses of the heart, spleen and brain are larger in legacy lines compared with modern broilers (13, 14, 21). In modern broilers, this likely is responsible for increased incidence of cardiomyopathy (22, 23) and immune deficiencies (10). The reduced brain mass in broilers could contribute to the behavioral differences seen in modern broilers compared with other lines. These include a reduction in movement, along with reduced fear and risk aversion in comparison with legacy or layer lines (24) In addition, a myopathy called Wooden Breast disease has arisen during the selection process (25-27). This myopathy is apparently due to selection for rapid growth and is characterized by necrosis, fibrosis, and immune cells infiltration that results in a product that is unappealing to consumers (28-31). This disease results in significant losses to the poultry industry as the meat is condemned at processing (32).

In prior work, we compared the breast muscle transcriptome of 6-day old (D6) modern and legacy lines (16). That comparison identified differently expressed genes affecting growth factors, lipid metabolism and the pentose phosphate pathway. In particular, the modern transcriptome indicated elevated expression of Insulin Like Growth Factor 1 (IGF1) which stimulates muscle hypertrophy combined with reduced expression of Myostatin (MSTN), an inhibitor of skeletal muscle hyperplasia (33). These studies suggest that human directed evolution increased growth factor stimulation in the modern breast muscle. Furthermore, elevated expression of genes involved in the pentose phosphate pathway and lipid metabolism would provide necessary intermediates and energy required for rapid growth.

Here we report a comparative study between modern Ross 708 and legacy UIUC breast muscle proteomes. The results support and extend the conclusions of our prior studies, providing further insight into the impact of selection that yielded modern broilers (14, 16). The data also have implications for the negative effects of intense selection on the broiler chicken along with the emergence of Wooden Breast disease (25, 28, 29, 34, 35)

## Materials and Methods

### Animal care and sample collection

Animal raising, handling and sample collection methods were approved by the Committee on the Ethics of Animal Experiments of the University of Delaware (Permit Number: 2703-12-10). Six male UIUC and six male Ross 708 breast muscle samples were collected at day 6 post-hatch, frozen in liquid nitrogen and stored at -80°C until processed for proteome analysis.

### Proteomic analysis

For each of the muscle samples, six technical replicates were analyzed by mass spectrometry. In each case 50 mg of each muscle sample was subject to differential detergent fractionation and 20 μg of each fraction was trypsin digestion as previously described (36, 37). Following digestion, each fraction was desalted using a peptide macrotrap (Michrom BioResources) according to the manufacturer’s instructions. After desalting, each fraction was further purified using a strong cation exchange macrotrap to remove any residual detergent, which could interfere with the mass spectrometry. Fractions were dried and resuspended in 10 μl of 2% acetonitrile, 0.1% formic acid and transferred to low retention vials in preparation for separation using reverse phase liquid chromatography.

An Ultimate 3000 (Dionex) high performance liquid chromatography system coupled with an LTQ Velos Pro (Thermo) mass spectrometer were for peptide separation and mass spectrum acquisition. The U3000 was operated at a flow rate of 333 nl per minute and equipped with a 75 μm x 10 cm fused silica column packed with Halo C18 reverse phase material (Mac-Mod Analytical). Each peptide sample was separated using a 4 h gradient from 2% to 50% acetonitrile with 0.1% formic acid as a proton source. The column was located on a Nanospray Flex Ion Source (Thermo) and connected directly to a silica Nanospray emitter to minimize peak broadening. High voltage was applied using a stainless-steel junction between the column and the emitter. Scan parameters for the LTQ Velos Pro were one MS scan followed by 10 MS/MS scans of the 5 most intense peaks. MS/MS scans were performed in pairs, one using collision induced dissociation (CID) and the other using higher-energy collisional dissociation (HCD). Dynamic exclusion was enabled with a mass exclusion time of 3 min and a repeat count of 1 within 30 sec of initial m/z measurement.

### Protein identification

Spectrum matching programs X!tandem (38) and OMSSA (39) were used via the University of Arizona High Throughput Computing Center. Raw spectra were converted to MGF format for analysis using the MSConvert tool from the ProteoWizard software suite (40). X!tandem was run with 12 threads, precursor and fragment tolerance of 0.2 Da, and up to two missed tryptic cleavages. Variable modifications used in the searches were: single and double oxidation of Methionine, carbamidomethylation of Cysteine, and phosphorylation of Tyrosine, Threonine, and Serine. OMSSA was run with 12 threads, precursor and fragment tolerance of 0.2 Da, up to two missed tryptic cleavages, and set to XML output format. A custom Perl script was used to parse XML search results from both X!tandem and OMSSA. Peptides with e-values ≤ 0.05 were accepted and single spectrum identifications were rejected unless they were identified by both search engines. To verify data set quality, decoy searches were performed in the exact manner as before, but with a randomized version of the protein databases. False discovery rates ranged from 0.9% to 2.3% with an average of 1.7%.

### Identifying differentially expressed proteins

Differential expression of proteins between treatments was performed pairwise using peptide elution profiles as described by Wright et al (41). Precursor mass spectra were extracted from the raw data in MS1 format using the MSConvert software from the ProteoWizard toolset. Peptide precursor m/z values were extracted from the previously compiled protein identifications using Perl. Peptide intensities were summed for each protein on a per-replicate basis. Data were normalized based on the mode of each replicate rather than the mean to minimize the effect of extreme values. A resampling analysis was performed for each pairwise comparison. Proteins were considered to be differentially expressed if the difference in means between conditions resulted in a P-value ≤ 0.05.

### Data deposition

Transcript data discussed in this publication have been deposited in NCBI’s Gene Expression Omnibus (GEO) and are accessible through GEO Series accession number GSE65217. Red Jungle Fowl proteomics data is available from ProteomeXchange (PXD005288). Modern Ross 708 and legacy UIUC proteomics data from day 6 muscle have also been deposited to ProteomeXchange (XXXX and XXXX, respectively).

## Results

### Proteomic Results

In the comparison between six male day 6 modern Ross 708 and six male day 6 legacy UIUC breast muscle, a total of 222 differentially expressed proteins were detected (p-value <0.05) with 173 enriched in modern samples and 49 enriched in legacy samples. Of the 222 differentially expressed proteins, 130 (57%) exhibited the same direction of enrichment as shown in an earlier transcriptome study (16) (Supporting Information Table S1).

### Gene Ontology

(GO) (42). Muscle from legacy birds were enriched for GO terms relevant to myofibers and energy production in striated muscle, along with vesicle transport (Fig. 1). Muscle from modern birds were enriched for terms including stress response, myofibers, translation, energy production, metabolism, vesicle transport and innate immunity.

**Figure 1.**
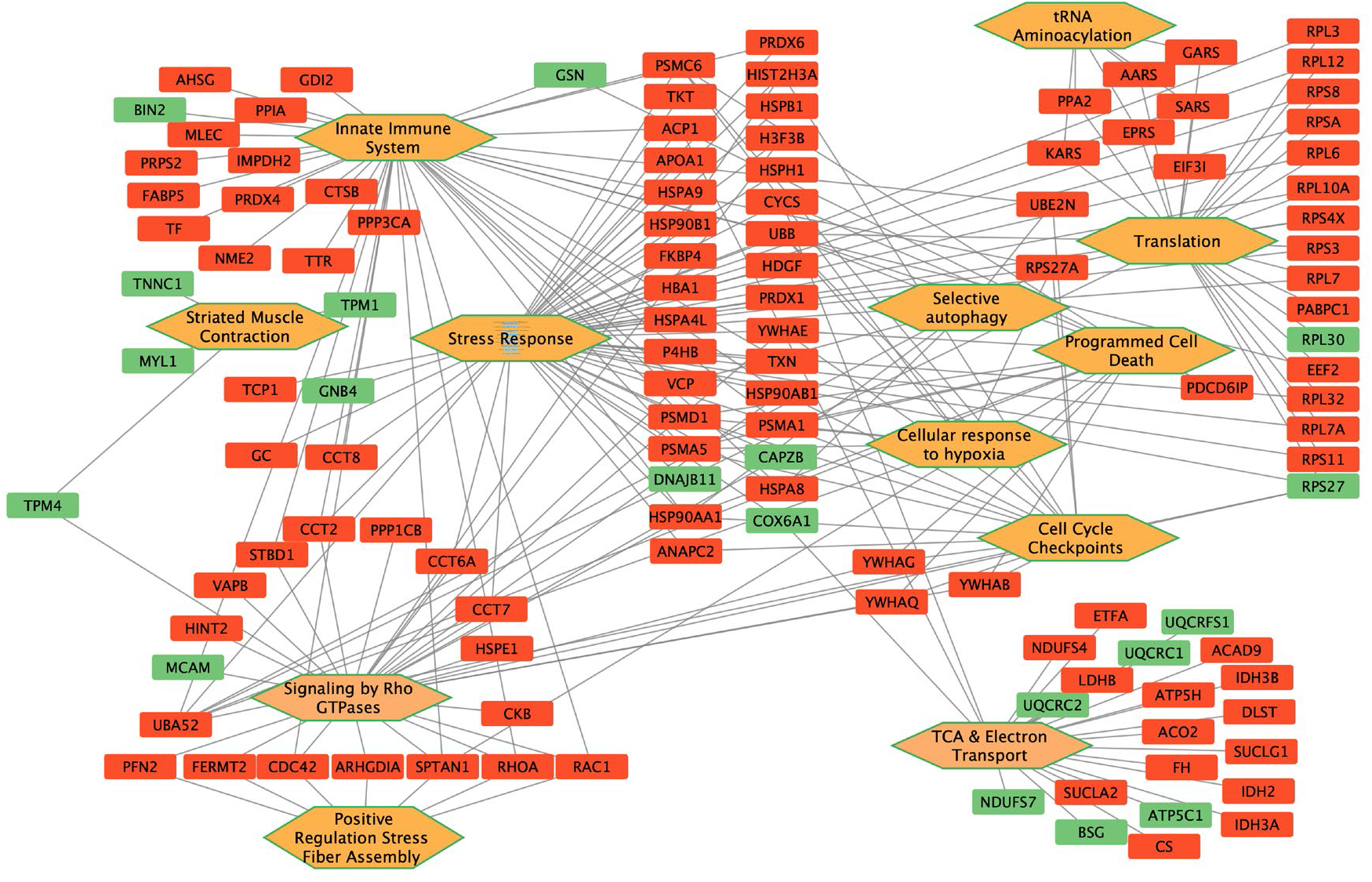
Functional enrichment in differentially expressed muscle. Orange hexagons refer to GO terms and rectangles refer to proteins differentially enriched between modern Ross 708 (Red) and legacy UIUC (Green) day 6 muscle.

**Figure 1**. Gene Ontology results. Orange hexagons refer to GO terms and rectangles refer to proteins differentially enriched between modern (Red) and legacy (Green) chicken muscle.

### Stress Response

In the comparison between modern and legacy muscle, 49 differentially regulated proteins were part of the stress response. Five stress response proteins were elevated in the legacy breast muscle including cytochrome c oxidase subunit 6A1 (COX6A1), capping actin protein (CAPZB), ribosomal protein S27 (RPS27), G protein subunit beta 4 (GNB4) and DnaJ heat shock protein family Hsp40 member B11 (DNAJB11). 44 stress-associated proteins were elevated in the modern chicken muscle compared to the legacy samples. These function in a variety of stress related processes including acting as chaperones and cochaperones, responding to genotoxic stress, redox regulation, and protein degradation. Among these were four proteasome subunits that likely offset the increased burden of misfolded proteins due to the large amount of protein synthesis required for muscle hypertrophy. Nine heat shock proteins (HSPs) were also elevated in muscle from the modern line, all of which function as chaperones controlling the proper folding of client proteins. Five members of the TCP complex that acts as an ATP dependent chaperone controlling actin folding were enriched in the modern samples. (5, 6). Four isomerases responsible for controlling disulfide or proline conformation and three proteins that control redox stress were also elevated in the modern line.

### Myofiber

Gene expression in legacy muscle showed enrichment for several myofibril proteins including Gelsolin (GSN), Troponin C (TNCC), Tropomyosin (TMP1 and TMP4), Myosin Light Chain 1 (MYL1), Myomesin 2 (MYOM2), Cytoskeletal Keratins (KRT5 and KRT7), and Fragile X Mental Retardation Syndrome-Related Protein (FXR1). CAPZB, GSN, TNCC, TPM1 and TPM4 regulate the dynamics of actin filament assembly. MYL1 is a non-regulatory light chain that interacts with actin in generating contraction. MYOM2 is a structural component that stabilizes the M band of muscle and KRT5 and KRT7 are intermediate filaments components. FXR1 controls mRNA transport and translation (43). Knockdown mutations of FXR1 in mice reduce limb musculature and result in early mortality (44) and recessive mutations in humans results in multi-minicore myopathy (45). The transcript encoding Bridging Integrator 1 (BIN1) is enriched in the modern line. BIN1 protein localizes to T-tubules (46) where it controls Ca^2+^ signaling (47). Mutations in BIN1 cause centronuclear myopathy, which causes muscle weakness and atrophy (48-50).

### Translation

The most striking contrast between modern broilers such as Ross 708 chickens and earlier breeds is the difference in both normalized and total breast muscle yield. Some of the increase in normalized breast muscle of modern broilers can be attributed to hypertrophy due to increased protein synthesis (51, 52). This is consistent with the enrichment in the Ross 708 birds for protein translation initiation factors and ribosomal structural proteins (Supplemental Table S1 and Fig. 1). In addition to ribosomal proteins, five enzymes encoding tRNA ligases were enriched in Ross 708 breast muscle including: lysine–tRNA ligase (KARS), serine–tRNA ligase (SARS), glycine–tRNA ligase (GARS), alanine–tRNA ligase (AARS) and bifunctional glutamate/proline– tRNA ligase (EPRS). These would support the elevated protein translation required for the muscle hypertrophy seen in the modern line.

### Glycolysis

Muscle from the UIUC legacy line is enriched for two proteins involved in glycolysis: Hexose Kinase (HK) and Phosphofructokinase (PFKM) (Fig. 2). HK catalyzes the first reaction of glycolysis by phosphorylating glucose while PFKM drives the commitment step to glycolysis. Elevated levels of these glycolytic enzymes are consistent with the fast twitch nature of breast muscle from wild chickens, which is driven by glycolysis.

**Figure 2.**
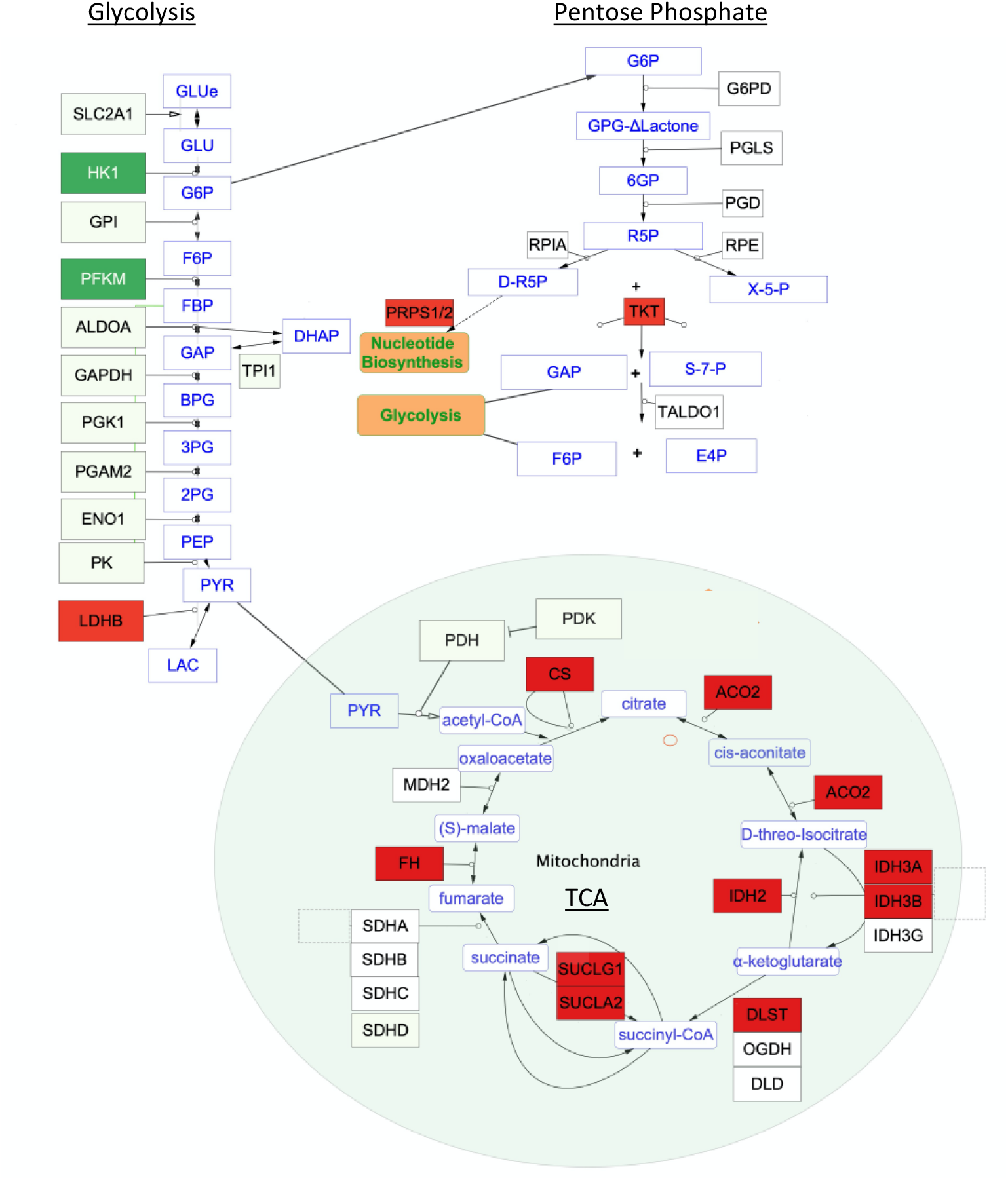
Glycolysis, Pentose Phosphate Pathway and TCA cycle. Red rectangles indicate enzymes enriched in modern Ross 708 chicken muscle while dark green indicates the enzymes enriched in legacy line muscle.

**Figure 2**. Glycolysis, Pentose Phosphate Pathway and TCA cycle. Red rectangles indicate enzymes enriched in the modern line while dark green indicates the enzymes enriched in the legacy line.

### Pentose Phosphate Pathway (PPP)

modern line breast muscle contained higher levels of the PPP enzymes transketolase (TKT) and phosphoribosyl pyrophosphate synthetase (PRPS1 and PRPS2). These enzymes function in the nonoxidative phase of PPP returning fructose 6 phosphate and glyceraldehyde 3 phosphate to glycolysis and providing ribose 5 phosphate as a precursor for purine and pyrimidine synthesis (Fig. 2).

### TCA cycle

The main pathway affecting energy production enriched in the modern line breast muscle was the TCA cycle (Fig. 2). Most enzymes directly involved in the TCA cycle were elevated in the modern line muscle, including: Citrate Synthase (CS), Aconitase (ACO2), Isocitrate Dehydrogenase (IDH, IDH3A. IDH3B), Dihydrolipoyllysine-residue Succinyltransferase (DLST, a component of α-Ketoglutarate Dehydrogenase), Succinyl-CoA ligase (SUCLA2 and SUCLG1) and Fumerase (FH). The components not elevated were succinate dehydrogenase (SDHA) and malate dehydrogenase (MDH). The enrichment for components of the TCA cycle indicates the central role of this pathway in the energy metabolism of the modern Ross 708 chickens compared with the legacy UIUC birds. In addition to providing coenzymes for oxidative phosphorylation, the TCA cycle provides intermediates for cataplerosis to be used as precursors for nucleotides, amino acids, and lipids. Anaplerosis could be supported by elevated levels of glutamic-oxaloacetic transaminase 2 (GOT2) that converts glutamate to α-ketoglutarate thus allowing for replenishing TCA cycle metabolites.

### Beta-Oxidation

Enzymes involved in fatty acid beta oxidation are enriched in the modern birds compared with the legacy line (Figure 3). One enriched enzyme is acyl-CoA dehydrogenase family member 9 (ACAD9), the rate limiting enzyme in the oxidation of fatty acyl CoA. ACAD9 is responsible for introducing a *trans* double bonds into palmitoyl-CoA and initiating the beta-oxidation of this common lipid. Also enriched is enoyl-CoA delta isomerase 1(ECI1) which is necessary for beta oxidation of unsaturated fatty acids. The transcript encoding hydroxyacyl-CoA dehydrogenase (HADH) is also elevated in the modern Ross 708 muscle. This enzyme acts repeatedly on lipids, sequentially removing two carbon units by oxidizing a 12-carbon fatty acid to acetoacetyl-CoA. Acetyl-CoA acyltransferase 2 (ACAT2), also enriched in Ross 708 breast muscle, oxidizes acetoacetyl-CoA to acetyl-CoA. The product of beta oxidation, acetyl-CoA, can then be metabolized via the TCA cycle to generate energy, or used in anabolic reactions to support rapid breast muscle growth.

**Figure 3.**
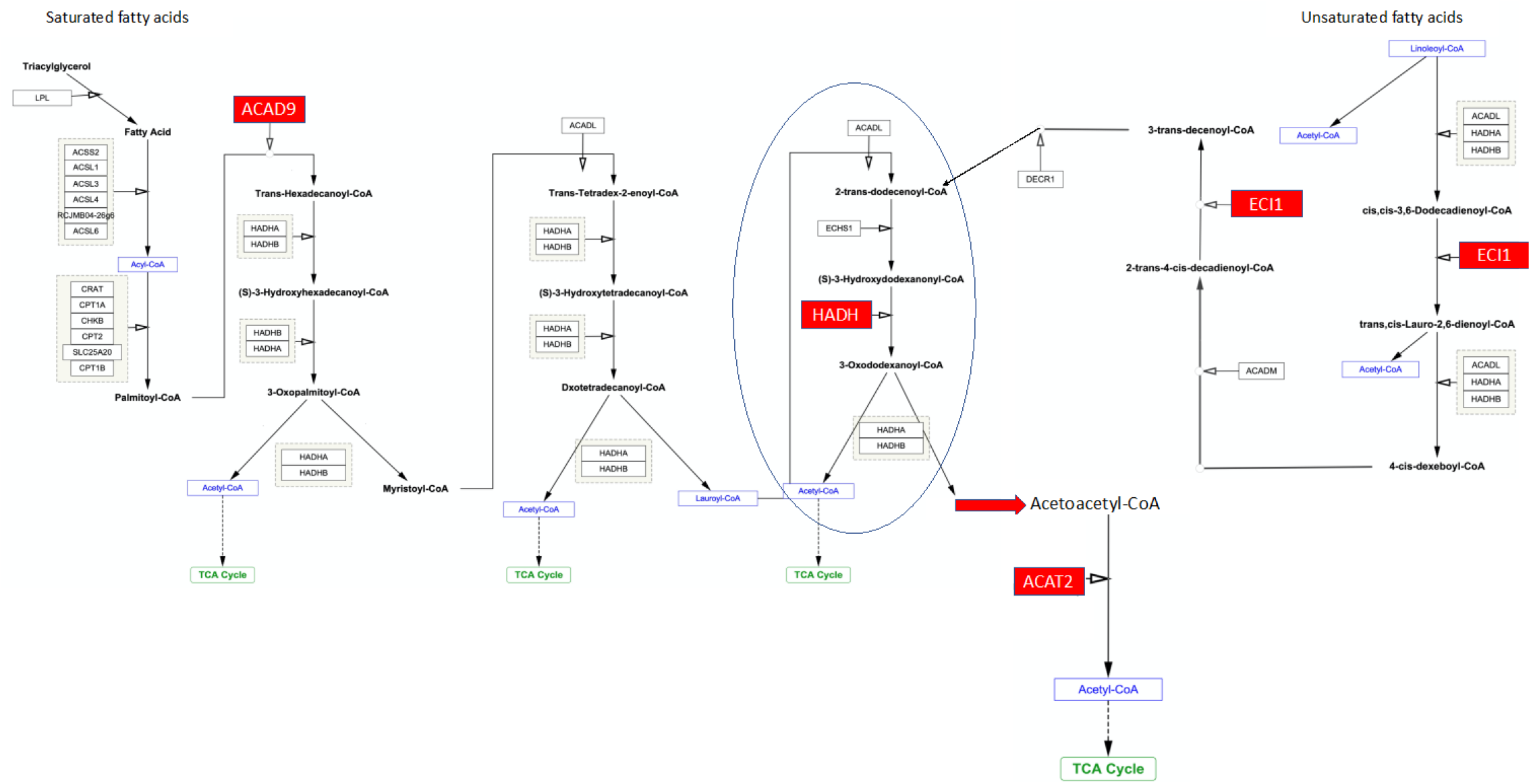
Beta-oxidation of Saturated Fatty Acids (SFA) and Unsaturated Fatty acids (USFA). Red indicates proteins that are enriched in modern Ross 708 muscle. The step indicated by the oval is repeated, sequentially removing Acetyl-CoA, until the 12 carbon Lauroyl-CoA is oxidized to the 4 carbon Acetoacetyl-CoA which is oxidized to Acetyl-CoA by ACAT2. Red rectangles indicate enzymes enriched in the modern Ross 708 muscle while green indicates the enzymes enriched in muscle from the legacy UIUC line.

**Fig. 3**. Beta-oxidation of Saturated Fatty Acids (SFA) and Unsaturated Fatty acids (USFA). Red indicates proteins that are enriched in the modern line. The step indicated by the oval is repeated, sequentially removing Acetyl-CoA, until the 12 carbon Lauroyl-CoA is oxidized to the 4 carbon Acetoacetyl-CoA which is oxidized to Acetyl-CoA by ACAT2. Red rectangles indicate enzymes enriched in the modern line while green indicates the enzymes enriched in the legacy line

### Amino Acid Metabolism

In addition to TCA enzymes affecting amino acid anaplerosis, modern skeletal muscle was enriched for enzymes affecting glycine, lysine, and methionine metabolism. Enzymes affecting glycine impact choline and creatine levels include aldehyde dehydrogenase 7 family member A1 (ALDH7A1), betaine—homocysteine S-methyltransferase (BHMT), dimethylglycine dehydrogenase (DMDGH) and glycine amidinotransferase (GATM). ALDH7A1, BHMT and DMDGH form a pathway in the conversion of choline to sarcosine that is found in high concentrations in skeletal muscle. Choline is a precursor to phosphatidylcholine, an important component of cellular membranes. GATM is part of the pathway that mediates the interconversion between creatine and glycine. Creatine is also found at high concentration in muscle as it is important for energy storage as creatine phosphate. Further supporting creatine phosphorylation is creatine kinase B (CKB), which is elevated in modern breast muscle compared with the legacy line tissue. Three enzymes enriched in the modern line are involved in S-adenosylmethionine (SAM) production from methionine. These include methionine adenosyltransferase (MAT1A), adenosylhomocysteinase (BHMT), and adenosylhomocysteinase (AHCY). SAM functions as a universal methyl donor in biological systems.

### Electron Transport Chain

Enriched in the legacy breast muscle were enzymes associated with the mitochondrial respirasome. These enzymes include NADH dehydrogenase (NDUFS7), cytochrome c reductases (UQCRFS1, UQCRC1, UQCRC2), cytochrome c oxidase (COX6A1) and ATP5C1, a component of the ATP synthase central stalk. Four mitochondrial components were found enriched in modern Ross 708 muscle: one subunit of the electron-transfer-flavoprotein (ETFA) along with inorganic pyrophosphatase (PPA2). One NADH dehydrogenase (NDUFS4) and one component of the ATPase stalk, ATP synthase, H+ transporting, mitochondrial F_o_ complex subunit D (ATP5H) were also elevated in the modern muscle.

### Vesicle and Protein Transport

The legacy line breast muscle was enriched for proteins affecting vesicle transport including: COPI coat complex subunit beta 2 (COPB2), transmembrane p24 trafficking protein-2 (TMED2), and Amphiphysin 1 (AMPH). COPB2 functions in retrograde transport from the Golgi to the ER and is involved in recognition of specific vesicle cargo proteins (53). The TMED proteins function in anterograde transport of vesicles from the ER to the Golgi apparatus and play roles in cargo selection (9). AMPH is enriched in neural tissue where it is involved in endocytosis through interaction with clathrin (54, 55). Some instances of the paraneoplastic Stiff-person Syndrome (SPS) (56, 57) are due to AMPH autoantibodies produced in breast and lung cancer patients. SPS is characterized by muscle spasms, rigidity and hypertrophy that arise from the effect of the autoantibody on nerve cells that control muscles (58). Modern Ross 708 breast muscle was enriched for vesicle regulatory components including: Cell Division Control Protein 42 (CDC42), RAB GDP Dissociation Inhibitor (GDI2), RAS-related C3 botulinum toxin substrate (RAC1) and transforming protein RHOA (RHOA).

### Innate Immunity

Gene Ontology analysis identified 39 proteins that function in innate immunity. Two, Gelsolin (GSN) and Bridging Integrator 2 (BIN2) were enriched in the legacy line breast muscle. Elevated expression of GSN is associated with a decreased response to a variety of inflammatory stimuli (59-64) and elevated BIN2 levels are associated with decreased phagocytic activity (65). The 26 proteins enriched in the modern Ross 708 breast muscle are associated with neutrophil degranulation. For example, CTSB is secreted during neutrophil degranulation and degrades collagen upon release into the extracellular space (66, 67). PRDX6 and RAC1, associate with and activate NADPH oxidase, a major source of reactive oxygen species (68-70). RAC1 (71), RHOA (72) and WD repeat domain 1 (WDR1) (73) participate in the polarization that drives neutrophil migration. Birds do not have cells named neutrophils, but neutrophil function is carried out by heterophils in avian species (74). It is reasonable to hypothesize that the innate immunity proteins are present in heterophils within the breast muscle.

## Discussion

The differences between modern broilers (Ross 708) and legacy lines (UIUC), arose from human directed evolution selecting for the broiler’s rapid growth, improved feed efficiency and increased breast muscle mass. Rapid growth is evidenced in the time from hatch to market. For legacy birds grown for market, that would typically take 16 weeks, while with modern birds it takes 7 weeks. The improved feed efficiency arose in part from selection lengthening the absorptive segments of the small intestine combined with earlier maturation of the liver (14). Selection for increased breast muscle has generated modern birds with more than twice the breast muscle mass of legacy lines. The proteomic results presented here indicate that selection caused metabolic reprogramming that supports the excessive breast muscle growth seen in the modern lines.

In chickens, skeletal muscle hyperplasia is thought to occur prior to hatch, and the increased size of the post-hatch muscle is largely due to hypertrophy (75). Hypertrophy is driven by controlling the balance between protein synthesis and degradation (51, 52). The enrichment of 12 ribosomal subunit proteins, initiation factors EIF3I and EIF4A4, elongation factor EEF2, along with five tRNA ligases likely play an important role in the increased level of muscle hypertrophy modern lines. Elevated protein synthesis could also cause the increase in stress response evidenced in the modern birds. The stress proteins function in a variety of processes including serving as chaperones or cochaperones, or in modulating genotoxic stress, redox regulation, and protein degradation.

The legacy line birds exhibited enrichment for HK1 and PFKM enzymes, which drive the first phase of glycolysis. PFKM catalyzes the rate limiting step of glycolysis, and its enrichment would direct Glucose 6-Phosphate down the glycolytic pathway. This is consistent with the typical fast-twitch breast muscle fibers that are seen in birds that exhibit brief episodes of flight. In contrast, modern birds are enriched in one protein (TKT) that supports the nonoxidative part of the pentose phosphate pathway (PPP). The non-oxidative portion of the PPP provides precursors for nucleotide synthesis and feeds glucose metabolites back into glycolysis. Modern Ross 708 birds also express higher levels of Phosphoribosyl Pyrophosphate Synthetase, which direct Ribose-5-Phosphate to nucleotide production. The elevated level of LDHB in the Ross 708 birds is expected to drive the lactate:pyruvate equilibrium towards pyruvate. This would retain pyruvate for further metabolism by the TCA cycle and limit the release of lactate for energy production by other tissues (76). Furthermore, glucose consumption supporting the modern broiler’s breast muscle has ramifications for brain development. As glucose is the main energy source for the brain, diversion of this nutrient to the breast muscle may cause the reduced brain growth seen in modern broilers compared with legacy chicken lines or jungle fowl (13).

The elevated levels of multiple enzymes of the TCA cycle in the Ross 708 breast muscle allows this pathway to meet demands of breast muscle hypertrophy. This is supported by studies in other species implicating elevated TCA cycle activity in hypertrophy. For example, enrichment of TCA metabolites was noted in a KLF10 mouse knockout model of soleus muscle hypertrophy (77) with similar results seen in aerobic exercise induced hypertrophy in humans (78, 79). Furthermore, resistance training induced hypertrophy in humans increases the activity of citrate synthase, the gateway enzyme to the TCA cycle (80).

Several enzymes involved in lipid beta-oxidation are also elevated in birds from the modern line. Elevated expression of ACAD9 is particularly informative as this is the rate-limiting enzyme controlling lipid oxidation and elevated HADH activity also plays a role in skeletal muscle hypertrophy (80). Increased ACAD9, HADH, and other enzymes involved in lipid oxidation indicates that modern Ross 708 birds are using lipid metabolism, in addition to the pentose phosphate and TCA pathways, to provide resources supporting expansion of the breast muscle.

Elevated lipid beta-oxidation in the breast muscle may have ramifications for morphometric changes seen in the growth of modern broilers. For example, the reduced normalized heart mass in the modern Ross 708 line compared with legacy UIUC birds could cause the cardiomyopathy seen in modern broilers. Also, if normalized spleen mass is viewed as a proxy for immune functions, the morphometric data indicates that immune function is significantly lower in modern lines compared with birds from the legacy line. Metabolically this may arise from the elevated lipid use in modern Ross 708 skeletal muscle. Cardiac muscle and the immune system use lipids as a major source of energy. Consequently, increased competition with skeletal muscle for lipids might inhibit heart growth and immune function seen in modern broilers. In addition to glucose the brain also readily uses ketone bodies, such as acetoacetate, to function. The elevated levels of ACAT2 in modern breast muscle may reduce the availability of ketone bodies for use by the brain.

These data support prior studies comparing the transcriptomes of 6-day old modern Ross 708 and legacy UIUC birds which concluded that the legacy line breast muscle was enriched for transcripts associated with glycolysis, while the transcriptome of the modern birds favored beta-oxidation (16). Additionally, TKT transcripts were also elevated in a study of birds with high feed efficiency compared with low feed efficiency chickens (26). Elevated levels of this enzyme seen in this proteome analysis provide further support for increased expression of these proteins improving feed efficiency.

While these samples were obtained at day 6 post-hatch, there are already differences between the modern and legacy lines that have implications for the development of Wooden Breast Disease (28). The elevated levels of stress proteins seen in breast muscle from the modern line provide compelling evidence that this tissue is undergoing a variety of stresses, likely due to its rapid growth. Oxidative stress is thought to be a major contributor to the development of this disease (28, 81, 82) and five of the stress responsive proteins detected in this study, PIT54 (83, 84), Peroxiredoxin 1, 4 and 6 and Thioredoxin play important roles in regulating oxidative stress. Gene Ontology analysis of our data also detected proteins associated with neutrophils in the mammalian immune system. Neutrophils are one of the earliest responders to inflammation and in birds the role of these phagocytic cells is filled by heterophils (74). Histological examination of birds prone to Wooden Breast Myopathy revealed heterophilic infiltration in the pectoral muscle of D14 chickens and this is thought to be an early sign of disease development (25). Furthermore, lipid metabolism has been shown to be altered in rapidly growing broilers that develop Wooden Breast Disease (85). Taken together, the proteomic data suggests that Wooden Breast Disease starts to develop well before the disease is visually or palpably evident.

## Conclusions

This study provides insight into the proteomic response to human directed selection for broiler production traits. Elevated levels of proteins involved in translation support the increased muscle hypertrophy seen in modern broilers. Changes in energy production pathways including increased fatty acid oxidation, diversion of glucose into the pentose phosphate pathway, retention of pyruvate through the action of LDHB and elevated TCA enzymes, provide the resources necessary for continued muscle growth of modern broilers. In turn, the activity of these pathways in the breast muscle are likely preventing nutrients from reaching other organs, thus leading to the health deficiencies and behavioral changes seen in modern broilers compared with legacy lines. Also, elevation of inflammatory response proteins in day 6 modern line muscle have implications for the rise of myopathies during broiler selection. The data suggest that Wooden Breast Disease may have its origins early in the post-hatch growth period. An important future direction will be to examine the proteomes of breast muscle tissue from these two lines after day 6. In addition, the identity of these differentially regulated proteins can be used to generate testable hypotheses regarding the regulatory mechanisms that orchestrate the changes that have been introduced to the broiler chicken by human selection.

## Acknowledgments

This projected was supported by the Agriculture and Food Research Institute competitive grant (2011-67003-30228) from the United States Department of Agriculture National Institute of Food and Agriculture

**Supporting Information Table S1** Differentially expressed proteins. Symbols with asterisks are ones that transcriptome data (16) supported proteome data.

