## Supplemental Table 1 for "Proteomic insight into human directed evolution of the domesticated chicken *Gallus gallus*"

Supporting Table S1

| **Name** | **Symbol** | **Enriched** | **P-value** |
| --- | --- | --- | --- |
| alpha-2-macroglobulin-like 4 | A2ML4 | Ross708 | 0.0215 |
| alanyl-tRNA synthetase 2, mitochondrial | AARS* | Ross708 | 0.0246 |
| acetyl-CoA acyltransferase 2 | ACAA2* | Ross708 | 0.0003 |
| acyl-CoA dehydrogenase family member 9 | ACAD9* | Ross708 | 0.0128 |
| acetyl-CoA acetyltransferase 2 | ACAT2* | Ross708 | 0.0048 |
| aconitase 2 | ACO2* | Ross708 | 0.0006 |
| acid phosphatase 1 | ACP1* | Ross708 | 0.0079 |
| adenosine kinase | ADK* | Ross708 | 0.0143 |
| adenosylhomocysteinase | AHCY* | Ross708 | 0.0005 |
| alpha 2-HS glycoprotein | AHSG | Ross708 | 0.0316 |
| aldehyde dehydrogenase 1 family member L2 | ALDH1L2* | Ross708 | 0.0095 |
| aldehyde dehydrogenase 6 family member A1 | ALDH6A1* | Ross708 | 0.0137 |
| aldehyde dehydrogenase 7 family member A1 | ALDH7A1 | Ross708 | 0.0304 |
| amphiphysin | AMPH* | UIUC | 0.0294 |
| anaphase promoting complex subunit 2 | ANAPC2* | Ross708 | 0.0098 |
| acidic nuclear phosphoprotein 32 family member A | ANP32A | UIUC | 0.0128 |
| acidic nuclear phosphoprotein 32 family member B | ANP32B | UIUC | 0.0049 |
| acidic nuclear phosphoprotein 32 family member E | ANP32E | UIUC | 0.0012 |
| annexin A6 | ANXA6* | Ross708 | 0.0009 |
| apolipoprotein A1 | APOA1* | Ross708 | 0.0049 |
| Rho GDP dissociation inhibitor alpha | ARHGDIA* | Ross708 | 0.0160 |
| 5-aminoimidazole-4-carboxamide ribonucleotide formyltransferase/IMP cyclohydrolase | ATIC* | Ross708 | <.0001 |
| ATPase Na+/K+ transporting family member beta 4 | ATP1B4* | UIUC | 0.0045 |
| ATP synthase, H+ transporting, mitochondrial F1 complex, gamma polypeptide 1 | ATP5C1 | UIUC | 0.0079 |
| ATP synthase, H+ transporting, mitochondrial F0 complex, subunit D | ATP5H* | Ross708 | 0.0466 |
| betaine--homocysteine S-methyltransferase | BHMT* | Ross708 | 0.0004 |
| bridging integrator 1 | BIN1* | Ross708 | 0.0428 |
| bridging integrator 2 | BIN2 | UIUC | 0.0263 |
| bleomycin hydrolase | BLMH* | Ross708 | 0.0023 |
| bisphosphoglycerate mutase | BPGM* | UIUC | 0.0419 |
| basigin (Ok blood group) | BSG | UIUC | 0.0135 |
| basic transcription factor 3 | BTF3 | Ross708 | 0.0006 |
| calpain 3 | CAPN3 | Ross708 | 0.0014 |
| capping actin protein of muscle Z-line subunit beta | CAPZB | UIUC | 0.0350 |
| carbonyl reductase 1 | CBR1* | Ross708 | 0.0479 |
| coiled-coil domain containing 47 | CCDC47* | UIUC | 0.0126 |
| chaperonin containing TCP1 subunit 2 | CCT2* | Ross708 | 0.0036 |
| chaperonin containing TCP1 subunit 6A | CCT6A* | Ross708 | 0.0239 |
| chaperonin containing TCP1 subunit 7 | CCT7* | Ross708 | 0.0151 |
| chaperonin containing TCP1 subunit 8 | CCT8* | Ross708 | 0.0114 |
| cell division cycle 42 | CDC42* | Ross708 | 0.0085 |
| cofilin 2 | CFL2* | Ross708 | <.0001 |
| coiled-coil-helix-coiled-coil-helix domain containing 3 | CHCHD3* | Ross708 | 0.0469 |
| creatine kinase B | CKB* | Ross708 | 0.0400 |
| carnosine dipeptidase 2 | CNDP2* | Ross708 | 0.0178 |
| collagen type VI alpha 1 chain | COL6A1* | Ross708 | 0.0440 |
| collagen type XV alpha 1 chain | COL15A1 | UIUC | 0.0396 |
| COPI coat complex subunit beta 2 | COPB2 | UIUC | 0.0298 |
| cytochrome c oxidase subunit 6A1, mitochondrial | COX6A1 | UIUC | 0.0076 |
| citrate synthase | CS | Ross708 | 0.0047 |
| cathepsin B | CTSB | Ross708 | 0.0394 |
| cullin 5 | CUL5 | Ross708 | 0.0388 |
| cytochrome c, somatic | CYCS | Ross708 | 0.0254 |
| dihydrolipoamide S-succinyltransferase | DLST* | Ross708 | 0.0226 |
| dimethylglycine dehydrogenase | DMGDH* | Ross708 | 0.0061 |
| DnaJ heat shock protein family (Hsp40) member B11 | DNAJB11 | UIUC | 0.0302 |
| destrin, actin depolymerizing factor | DSTN* | Ross708 | 0.0048 |
| enoyl CoA hydratase 1, peroxisomal | ECH1 | Ross708 | 0.0260 |
| enoyl-CoA delta isomerase 1 | ECI1* | Ross708 | 0.0429 |
| eukaryotic translation elongation factor 2 | EEF2* | Ross708 | 0.0027 |
| eukaryotic translation initiation factor 3 subunit I | EIF3I* | Ross708 | 0.0492 |
| eukaryotic translation initiation factor 4A3 | EIF4A3* | Ross708 | 0.0493 |
| ER membrane protein complex subunit 1 | EMC1 | UIUC | 0.0204 |
| ectonucleoside triphosphate diphosphohydrolase 2 | ENTPD2 | UIUC | 0.0310 |
| glutamyl-prolyl-tRNA synthetase 1 | EPRS* | Ross708 | 0.0006 |
| electron transfer flavoprotein subunit alpha | ETFA* | Ross708 | 0.0344 |
| fatty acid binding protein 5 | FABP5* | Ross708 | 0.0041 |
| fermitin family member 2 | FERMT2 | Ross708 | 0.0378 |
| fumarate hydratase | FH | Ross708 | 0.0040 |
| FKBP prolyl isomerase family member 1C | FKBP1C | Ross708 | 0.0038 |
| FKBP prolyl isomerase 4 | FKBP4* | Ross708 | 0.0028 |
| FMR1 autosomal homolog 1 | FXR1 | UIUC | 0.0155 |
| glycyl-tRNA synthetase 1 | GARS* | Ross708 | 0.0089 |
| glycine amidinotransferase | GATM* | Ross708 | 0.0017 |
| GC vitamin D binding protein | GC | Ross708 | 0.0091 |
| GDP dissociation inhibitor 2 | GDI2* | Ross708 | 0.0069 |
| G protein subunit beta 4 | GNB4 | UIUC | 0.0260 |
| glutamic-oxaloacetic transaminase 2 | GOT2 | Ross708 | 0.0377 |
| gelsolin | GSN* | UIUC | 0.0471 |
| H3 histone, family 3B (H3.3B) | H3F3B | Ross708 | 0.0135 |
| H3 histone, family 3C | H3F3C* | Ross708 | 0.0135 |
| hydroxyacyl-CoA dehydrogenase | HADH* | Ross708 | 0.0272 |
| hemoglobin subunit alpha 1 | HBA1 | Ross708 | 0.0173 |
| hemoglobin subunit mu | HBM | Ross708 | 0.0032 |
| heparin binding growth factor | HDGF | Ross708 | 0.0260 |
| 3-hydroxyisobutyrate dehydrogenase | HIBADH* | Ross708 | 0.0044 |
| histidine triad nucleotide binding protein 2 | HINT2 | Ross708 | 0.0321 |
| histone cluster 1, H1.01 | HIST1H101 | Ross708 | 0.0349 |
| histone cluster 1, H1.03 | HIST1H103 | Ross708 | 0.0386 |
| histone cluster 1, H1.10 | HIST1H110 | Ross708 | 0.0457 |
| histone cluster 1, H1.11 Like | HIST1H111L | Ross708 | 0.0294 |
| histone cluster 2 H3 | HIST2H3A | Ross708 | 0.0135 |
| hexokinase 1 | HK1* | UIUC | 0.0314 |
| heterogeneous nuclear ribonucleoprotein A1 | HNRNPA1 | UIUC | 0.0012 |
| heterogeneous nuclear ribonucleoprotein D like | HNRNPDL | UIUC | 0.0064 |
| heterogeneous nuclear ribonucleoprotein H2 | HNRNPH2 | UIUC | 0.0167 |
| heterogeneous nuclear ribonucleoprotein R | HNRNPR | UIUC | 0.0338 |
| heat shock protein 90 alpha family class A member 1 | HSP90AA1 | Ross708 | 0.0015 |
| heat shock protein 90 alpha family class B member 1 | HSP90AB1* | Ross708 | 0.0034 |
| heat shock protein 90 beta family member 1 | HSP90B1* | Ross708 | 0.0002 |
| heat shock protein family A (Hsp70) member 4 like | HSPA4L* | Ross708 | 0.0397 |
| heat shock protein family A (Hsp70) member 8 | HSPA8 | Ross708 | 0.0465 |
| heat shock protein family A (Hsp70) member 9 | HSPA9* | Ross708 | 0.0136 |
| heat shock protein family B (small) member 1 | HSPB1* | Ross708 | <.0001 |
| heat shock protein family D (Hsp60) member 1 | HSPD1* | Ross708 | 0.0003 |
| heat shock protein family E (Hsp10) member 1 | HSPE1* | Ross708 | 0.0004 |
| heat shock protein family H (Hsp110) member 1 | HSPH1* | Ross708 | 0.0371 |
| isocitrate dehydrogenase 2 | IDH2 | Ross708 | 0.0006 |
| isocitrate dehydrogenase 3 catalytic subunit alpha | IDH3A | Ross708 | 0.0236 |
| isocitrate dehydrogenase 3 | IDH3B | Ross708 | 0.0238 |
| inosine monophosphate dehydrogenase 2 | IMPDH2* | Ross708 | 0.0009 |
| integrin subunit alpha 6 | ITGA6* | Ross708 | 0.0188 |
| lysyl-tRNA synthetase 1 | KARS* | Ross708 | 0.0094 |
| keratin 5 | KRT5 | UIUC | 0.0268 |
| keratin 7 | KRT7 | UIUC | 0.0289 |
| lactate dehydrogenase B | LDHB* | Ross708 | 0.0002 |
| lumican | LUM* | Ross708 | 0.0371 |
| microtubule associated protein tau | MAPT | Ross708 | 0.0480 |
| methionine adenosyltransferase 1A | MAT1A* | Ross708 | 0.0250 |
| melanoma cell adhesion molecule | MCAM | UIUC | 0.0259 |
| malectin | MLEC | Ross708 | 0.0356 |
| myosin light chain 1 | MYL1* | UIUC | 0.0245 |
| myomesin 2 | MYOM2 | UIUC | 0.0118 |
| nucleolin | NCL | UIUC | 0.0305 |
| NADH:ubiquinone oxidoreductase core subunit S4 | NDUFS4* | Ross708 | 0.0219 |
| NADH:ubiquinone oxidoreductase core subunit S7 | NDUFS7 | UIUC | 0.0409 |
| nidogen 2 | NID2 | UIUC | 0.0002 |
| NME/NM23 nucleoside diphosphate kinase 2 | NME2* | Ross708 | 0.0381 |
| nucleolar and coiled-body phosphoprotein 1 | NOLC1* | Ross708 | 0.0216 |
| aminopeptidase like 1 | NPEPL1 | UIUC | 0.0148 |
| nucleophosmin 1 | NPM1* | Ross708 | 0.0358 |
| ornithine aminotransferase | OAT | Ross708 | 0.0007 |
| oncomodulin 2 | OCM2 | Ross708 | 0.0272 |
| Obg like ATPase 1 | OLA1 | Ross708 | 0.0331 |
| 3-oxoacid CoA-transferase 1 | OXCT1* | Ross708 | 0.0226 |
| prolyl 4-hydroxylase subunit beta | P4HB* | Ross708 | 0.0214 |
| poly(A) binding protein cytoplasmic 1 | PABPC1* | Ross708 | 0.0005 |
| protein kinase C and casein kinase substrate in neurons 3 | PACSIN3* | Ross708 | 0.0157 |
| phosphoribosylaminoimidazole carboxylase and phosphoribosylaminoimidazolesuccinocarboxamide synthase | PAICS* | Ross708 | 0.0246 |
| PBX homeobox interacting protein 1 | PBXIP1 | Ross708 | 0.0420 |
| protein-L-isoaspartate (D-aspartate) O-methyltransferase | PCMT1 | Ross708 | 0.0161 |
| programmed cell death 6 interacting protein | PDCD6IP* | Ross708 | 0.0396 |
| protein disulfide isomerase family A member 3 | PDIA3* | Ross708 | 0.0004 |
| phosphofructokinase, muscle | PFKM | UIUC | 0.0224 |
| profilin 2 | PFN2* | Ross708 | 0.0073 |
| prohibitin 2 | PHB2* | Ross708 | 0.0356 |
| PIT54 protein | PIT54 | Ross708 | 0.0487 |
| inorganic pyrophosphatase 2 | PPA2* | Ross708 | 0.0062 |
| peptidylprolyl isomerase A | PPIA | Ross708 | 0.0013 |
| protein phosphatase 1 catalytic subunit beta | PPP1CB* | Ross708 | 0.0323 |
| protein phosphatase 3 catalytic subunit alpha | PPP3CA* | Ross708 | 0.0289 |
| peroxiredoxin 1 | PRDX1* | Ross708 | 0.0003 |
| peroxiredoxin 4 | PRDX4* | Ross708 | 0.0361 |
| peroxiredoxin 6 | PRDX6* | Ross708 | 0.0131 |
| prolyl endopeptidase | PREP* | Ross708 | 0.0412 |
| protein kinase cAMP-dependent type II regulatory subunit alpha | PRKAR2A* | Ross708 | 0.0246 |
| Proline synthetase | PROSC* | Ross708 | 0.0412 |
| phosphoribosyl pyrophosphate synthetase 1 | PRPS1 | Ross708 | 0.0102 |
| phosphoribosyl pyrophosphate synthetase 2 | PRPS2* | Ross708 | 0.0060 |
| proteasome 20S subunit alpha 1 | PSMA1* | Ross708 | 0.0025 |
| proteasome 20S subunit alpha 5 | PSMA5 | Ross708 | 0.0003 |
| proteasome 26S subunit, ATPase 6 | PSMC6* | Ross708 | 0.0360 |
| proteasome 26S subunit, non-ATPase 1 | PSMD1* | Ross708 | 0.0366 |
| Rac family small GTPase 1 | RAC1* | Ross708 | 0.0105 |
| retinol binding protein 4 A | RBP4A | Ross708 | 0.0357 |
| ras homolog family member A | RHOA* | Ross708 | 0.0003 |
| ribosomal protein L3 | RPL3* | Ross708 | 0.0167 |
| ribosomal protein L6 | RPL6* | Ross708 | 0.0271 |
| ribosomal protein L7 | RPL7* | Ross708 | 0.0114 |
| ribosomal protein L7a | RPL7A* | Ross708 | 0.0109 |
| ribosomal protein L10a | RPL10A* | Ross708 | 0.0176 |
| ribosomal protein L12 | RPL12* | Ross708 | 0.0283 |
| ribosomal protein L30 | RPL30 | UIUC | 0.0491 |
| ribosomal protein L32 | RPL32 | Ross708 | 0.0466 |
| ribosomal protein S3 | RPS3* | Ross708 | 0.0078 |
| ribosomal protein S4 X-linked | RPS4X* | Ross708 | 0.0245 |
| ribosomal protein S8 | RPS8* | Ross708 | 0.0139 |
| ribosomal protein S11 | RPS11* | Ross708 | 0.0208 |
| ribosomal protein S27 | RPS27 | UIUC | 0.0219 |
| ribosomal protein S27a | RPS27A* | Ross708 | 0.0035 |
| ribosomal protein SA | RPSA* | Ross708 | 0.0246 |
| seryl-tRNA synthetase 1 | SARS | Ross708 | 0.0274 |
| serpin family A member 4 | SERPINA4 | Ross708 | 0.0182 |
| SET and MYND domain containing 1 | SMYD1* | Ross708 | 0.0331 |
| syntrophin alpha 1 | SNTA1 | Ross708 | 0.0070 |
| striated muscle enriched protein kinase | SPEG* | Ross708 | 0.0297 |
| SPHK1 interactor, AKAP domain containing | SPHKAP* | Ross708 | 0.0207 |
| spectrin alpha, non-erythrocytic 1 | SPTAN1* | Ross708 | 0.0426 |
| serine and arginine rich splicing factor 6 | SRSF6 | UIUC | 0.0004 |
| starch binding domain 1 | STBD1 | Ross708 | 0.0400 |
| succinate-CoA ligase ADP-forming subunit beta | SUCLA2* | Ross708 | <.0001 |
| succinate-CoA ligase GDP/ADP-forming subunit alpha | SUCLG1 | Ross708 | 0.0270 |
| t-complex 1 | TCP1* | Ross708 | 0.0158 |
| transferrin | TF | Ross708 | 0.0032 |
| thimet oligopeptidase 1 | THOP1 | Ross708 | 0.0003 |
| transketolase | TKT* | Ross708 | 0.0306 |
| transmembrane p24 trafficking protein 2 | TMED2 | UIUC | 0.0269 |
| troponin C1, slow skeletal and cardiac type | TNNC1* | UIUC | 0.0317 |
| tropomyosin 1 | TPM1* | UIUC | 0.0388 |
| tropomyosin 4 | TPM4 | UIUC | 0.0436 |
| tripeptidyl peptidase 2 | TPP2* | Ross708 | 0.0016 |
| tumor protein, translationally controlled 1 | TPT1 | Ross708 | 0.0031 |
| transthyretin | TTR | Ross708 | 0.0286 |
| taxilin beta | TXLNB | UIUC | 0.0392 |
| thioredoxin | TXN* | Ross708 | 0.0104 |
| ubiquitin A-52 residue ribosomal protein fusion product 1 | UBA52 | Ross708 | 0.0356 |
| ubiquitin B | UBB* | Ross708 | 0.0334 |
| ubiquitin conjugating enzyme E2 L3 | UBE2L3* | Ross708 | 0.0367 |
| ubiquitin conjugating enzyme E2 N | UBE2N* | Ross708 | 0.0090 |
| cytochrome b-c1 complex subunit 1, mitochondrial | UQCRC1 | UIUC | <.0001 |
| cytochrome b-c1 complex subunit 2, mitochondrial | UQCRC2 | UIUC | 0.0004 |
| cytochrome b-c1 complex subunit Rieske, mitochondrial | UQCRFS1 | UIUC | 0.0159 |
| VAMP associated protein B and C | VAPB* | Ross708 | 0.0289 |
| valosin containing protein | VCP* | Ross708 | 0.0018 |
| voltage dependent anion channel 1 | VDAC1* | UIUC | 0.0080 |
| voltage dependent anion channel 2 | VDAC2 | UIUC | 0.0018 |
| voltage dependent anion channel 3 | VDAC3 | UIUC | 0.0470 |
| WD repeat domain 1 | WDR1* | Ross708 | 0.0010 |
| tyrosine 3-monooxygenase/tryptophan 5-monooxygenase activation protein B | YWHAB* | Ross708 | 0.0165 |
| tyrosine 3-monooxygenase/tryptophan 5-monooxygenase activation protein E | YWHAE* | Ross708 | 0.0361 |
| tyrosine 3-monooxygenase/tryptophan 5-monooxygenase activation protein G | YWHAG* | Ross708 | 0.0129 |
| tyrosine 3-monooxygenase/tryptophan 5-monooxygenase activation protein Q | YWHAQ* | Ross708 | 0.0320 |
